# Emotional states affect walking performance

**DOI:** 10.1101/2023.03.29.534813

**Authors:** Abhishesh Homagain, Kaylena A. Ehgoetz Martens

**Affiliations:** Department of Kinesiology and Health Sciences, Faculty of Health, University of Waterloo, Waterloo, Ontario, Canada

## Abstract

Gait is a large component and indicator of health in both young and older adults. Many factors affect gait including age, disease, and even mood disorders. Few studies have looked at the influence of emotional states on gait. This study aimed to investigate the influence of emotional states on walking performance to understand whether an emotional state may be an important factor to consider when evaluating gait. Thirty-six young adults were recruited (23F, 13M) and performed a neutral baseline condition of walking which included six passes of walking across an 8m walkway (a total of 48m of walking). Participants then completed 6 pseudo-randomized emotional state induction conditions while immersive 360-degree videos were used to induce the following emotional state conditions: happiness, excitement, sadness, fear, and anger. Participants viewed the emotion elicitation videos using a virtual reality head-mounted display (HMD), then rated their emotional state using self-assessment manikins and walked (without the HMD) over a pressure sensor walkway. One-way repeated measures ANOVA and pairwise comparisons were used to examine differences in gait parameters across the emotional conditions. Participants walked with significantly reduced step length and speed during the sadness condition compared to the other emotion conditions and the neutral condition. Furthermore, participants adjusted the timing of their walking during the sadness condition and walked with significantly increased step, stance, and swing times compared to other emotion conditions, but not the neutral condition. Step time was significantly reduced during the conditions of excitement and fear compared to the neutral condition. These findings show that in young healthy adults, emotions may impact variety of gait parameters involving pace and rhythm, however have little influence on gait variability and postural control. These results indicate that perhaps the emotions of sadness and excitement should be taken into account as potential confounds for future gait analysis.

## Introduction

Human gait is a complex behaviour that is dynamic and adaptable, achieved through bidirectional interactions between the motor system of the brain and other cortical and sub-cortical structures related to cognitive-emotional functions [1,2]. A growing body of research has focused on understanding the influence of emotion on gait. For example, several studies have found that depressed individuals demonstrate marked changes to spatiotemporal gait parameters including reduced velocity, stride length, double limb support, and increased gait variability compared to controls [3–6]. Likewise, similar reductions to gait speed and step length as well as increases in step length variability and step time variability were also noted in highly anxious Parkinson’s patients compared to low anxiety Parkinson’s patients and age matched controls [7]. Whilst evidence suggests that mood disorders may give rise to discriminative changes in gait, less work has examined whether similar changes to gait can be induced by alterations in emotional state.

Studies have shown that particular emotional states such as happiness and anger feature faster walking with an increased stride length, and increased step count. Whereas other emotional states such as sadness and fear feature a reduction in speed, stride length, and step count [8–10]. Much of this past work has focused on quantifying aspects of pace, rather than variability of gait even though recent research has emphasized the clinical relevance of gait variability [11,12]. Furthermore, it is also important to note that most of the previous studies used a sample of professional actors to quantify emotional differences in walking, which can bias or exaggerate effect size of the results given that skilled actors are highly trained to produce stereotyped expressions [8–10]. Thus, there remains a gap in understanding the influence of emotional states on gait behaviour. Given that gait assessments can serve as an important marker for health status, it is imperative to understand the changes in gait characteristics that arise from fluctuations in emotional states; as these changes can possibly confound the gait characteristics observed in both healthy and non-healthy populations, leading to potential false characterizations of gait characteristics.

The main objective of this study is to measure the effect of emotional states on gait characteristics in healthy young adults with no acting experience using a broader set of gait parameters. It was hypothesized that emotions such as happiness, excitement and anger would result in increased gait speed and step length compared to the neutral control emotion [8–10]. Consequently, emotions such as fear and sadness were expected to result in the opposite, showing decreased gait speed [1,8]. Finally, based on past work in clinical populations, it was also hypothesized that fear would lead to an increase in gait variability.

## Methods

Thirty-six (n = 36) healthy young adult participants from the University of Waterloo were recruited for this study (Table1). Exclusion criteria included any previous difficulty experienced with virtual reality (VR) such as nausea, lightheadedness, fatigue etc. Recent history (6 months prior) of physical injuries that impacted gait, use of assistive devices for walking, or clinical diagnosis of mood disorders were also part of the exclusion criteria. Any participants taking medication that may induce or attenuate emotions were also excluded from the study. G*Power3 (Version33.1, Universitat Dusseldorf, Dusseldorf, Germany) was used to determine sample size using the effect size reported by Halovic & Kroos [9]. Effect size from gait speed was used as the key dependent variable as it is a reoccurring parameter measured across many of the studies. Using an alpha error probability of 0.05, power of 0.8, and the reported effect size of 0.64, a sample size of 18 was calculated [9]. The study was approved by the University of Waterloo Research Ethics Board. All participants provided written informed consent to participate in the study prior to collection. A virtual reality simulator sickness questionnaire (VRSQ) was used pre and post collection to account for possible motion sickness due to the VR environment.

**Table 1.** Demographic Information.

| Demographic Information | N = 36 |
| --- | --- |
| Sex, N (%) | 27F (75%) \| 9M (25%) |
| Age (Mean, Range) | 24(20-32) yrs |
| Height (Mean, Range) | 166(152 – 185) cm |
| mDES -Positive Emotions (Mean, Range) | 2.64 (0-4) |
| mDES-Negative Emotions (Mean, Range) | 1.21 (0-4) |
| Pre VRSQ-Score (Mean, Range) | 3.70 (0 - 22.50) |
| Post VRSQ-Score (Mean, Range) | 7.55 (0 - 30) |

*Demographic information of the $n=36$ sample of recruited participants.*

### Experimental Procedure

After initial demographic, eligibility and consent information was collected, a modified differential emotional questionnaire was used to screen initial emotional states of participants. Participants were then outfitted with an HTC Vive head mounted display (HTC, USA). VR content was generated using Unity (Unity Technologies, SF, CA, USA) and immersive videos were embedded inside the VR environment to induce an array of emotional states using Unity. This study used videos from the repository of videos created by Li et al. [13] as it had been previously used in VR scenarios, specifically for emotional elicitation. Notably, research has shown that VR is a useful and effective tool to elicit emotion [14,15] due to the benefit of the immersive environments that VR can create [14,16].

Participants were given time to explore and navigate the virtual environment to familiarize themselves with the novel stimuli prior to collection. After familiarization was completed, participants performed three neutral walking trials. Walking trials involved 6 passes of walking across the 6m gait carpet resulting in approximately 40-60 total steps captured. To account for gait acceleration, participants started their walks 1m before the carpet and walked an additional 1m past the edge of the carpet on each pass, resulting in a total walk distance of 8m per pass (48m total). These number of passes was chosen ensure that enough steps would be recorded to get valid measurements gait parameters [17]. All walking trials occurred in the real-world while the VR was used only for emotion elicitation. The emotional blocks were always completed after the neutral walking trials. Video of the participants on each walking trial was recorded during collection but was only used to confirm results from the pressure sensor carpet and was not used in any subsequent analysis.

Five emotional states of happiness, excitement, sadness, fear, and anger were elicited in a pseudorandomized blocked design. The blocks started with watching a video, then completing a pre-walk self-assessment manikin (SAM – which evaluated arousal, valence and dominance), followed by the walking trial, and culminating in the post-walk SAM. This sequence was repeated for a second video (eliciting the same emotion). Thus, each emotional state block consisted of 2 emotional induction videos and 2 walking trials. The videos were played inside the virtual environment and the participants were seated during viewing. Participants were asked to describe what emotions they were feeling after each video to validate the emotion elicitation protocol. For walking, participants were instructed to walk using their own self-selected pace and reflect upon the video as they were walking. The SAM was used to collect self-reported data on participant’s current emotional state. This was further used to validate whether the protocol was successful in eliciting the proper emotions alongside their self-reported emotional state. Given the heterogeneity of emotional states induced with the videos, the emotional response on the SAM in combination with the self-reported emotion state was used to confirm correct emotional elicitation in order to include the trial for gait analysis. Small breaks of 5 mins were offered between each block (but not between the video watching to walking portion) to reduce carryover effects of a certain emotion.

Each block was pseudo-randomized for each participant, with the positive emotions of happiness and excitement always being shown before the negative emotions of sadness, anger, and fear. This was due to feedback from a pilot study where participants reported difficulty experiencing positive emotions after having felt negative emotions. Notably, this effect was not reported in the opposite direction (participants did not report issues feeling negative emotions following feeling positive emotions). Five emotional state blocks in total were completed and a post-collection simulator sickness questionnaire was completed to account for any issues with motion sickness caused by the VR environment.

### Gait Parameters

Spatiotemporal parameters of gait were measured using the Zeno™ Walkway (ProtoKinetics, LLC, Havertown, USA) gait carpet. The PKMAS software (ProtoKinetics, LLC, Havertown, USA) was used to process and export the gait data. During data export, the first and last step made by the participant was excluded to control for the effects of acceleration and deceleration. The 5-factor model of gait, created by Lord et al. was used for gait analysis as it contains clinically relevant parameters which have been largely unexplored in studies investigating emotion and gait (specifically gait variability) [11]. All measures of gait variability were calculated using the coefficient of variation (% CV). Table 2 shows the list of all examined gait parameters.

**Table 2.** Gait domains and parameters.

| Gait Domain | Gait Parameters |
| --- | --- |
| **Pace** | Mean Step Velocity |
|  | Mean Step Length |
|  | Step Time Variability |
|  | Swing Time Variability |
| **Rhythm** | Mean Step Time |
|  | Mean Swing Time |
|  | Mean Stance Time |
| **Variability** | Step Velocity Variability |
|  | Step Length Variability |
|  | Step Width Variability |
| **Postural Control** | Mean Step Width |

*Shows the list of specific proposed gait parameters intended to be examined. All measures of variability were determined using % coefficient of variation, calculated by the ratio of standard deviation divided by the sample mean.*

### Statistical Analysis

A one-way repeated measures ANOVA was conducted with the six emotional state conditions representing the within-condition factor for each of the gait parameters listed in Table 2. The assumptions of normality were violated (measured via Shapiro-Wilks test) in at least one condition in all of the gait parameters with the exception of mean step width. Thus, a non-parametric Friedman’s ANOVA was used when assessing differences in gait parameters within all conditions except for mean step width where a parametric one-way ANOVA was used. Durbin-Conover pairwise comparisons or the Student’s t-test were also conducted with Bonferroni’s corrections upon reaching significant result (p<0.05) with the Friedman’s ANOVA or the parametric equivalent, respectively. All statistical test were performed in R Studio with R version 4.2.2. The ggstatsplot package was also used to perform the pairwise comparisons and generate the box plots [18].

Whilst all 36 participants watched almost every video and completed all walking trials, not all participants reported feeling the emotion that was intended. In addition, some participants reported that they were uncomfortable with certain videos and opted out of participating in certain emotional blocks (mainly fear). This resulted in a mismatch between emotional conditions and walking trials assessed across participants, making pairwise and repeated measures analysis a challenge. To account for these individual differences in emotional responses, a sample greater than the formally calculated sample size was recruited and a subset of 23 out of the 36 participants that explicitly reported they felt *each* of the targeted emotions was used for analysis.

## Results

Table 1 shows that participants generally did not report feeling extremely positive or extremely negative emotions prior to collection as shown by the moderate positive and negative mDES emotion scores. Although there was a reported increase in the post-VRSQ scores compared to the pre-VRSQ, participants did not report that their walking behaviour was significantly altered by viewing the videos. Significant differences in gait were only found in the domains of pace and rhythm while gait outcomes from the domains of variability and postural control showed no significant results. Table 3 shows the summary of the mean (SD) of each gait parameter examined across each emotion condition alongside the resultant p value of the Friedman’s tests (with the exception of step width where a one-way parametric ANOVA was used).

**Table 3.**
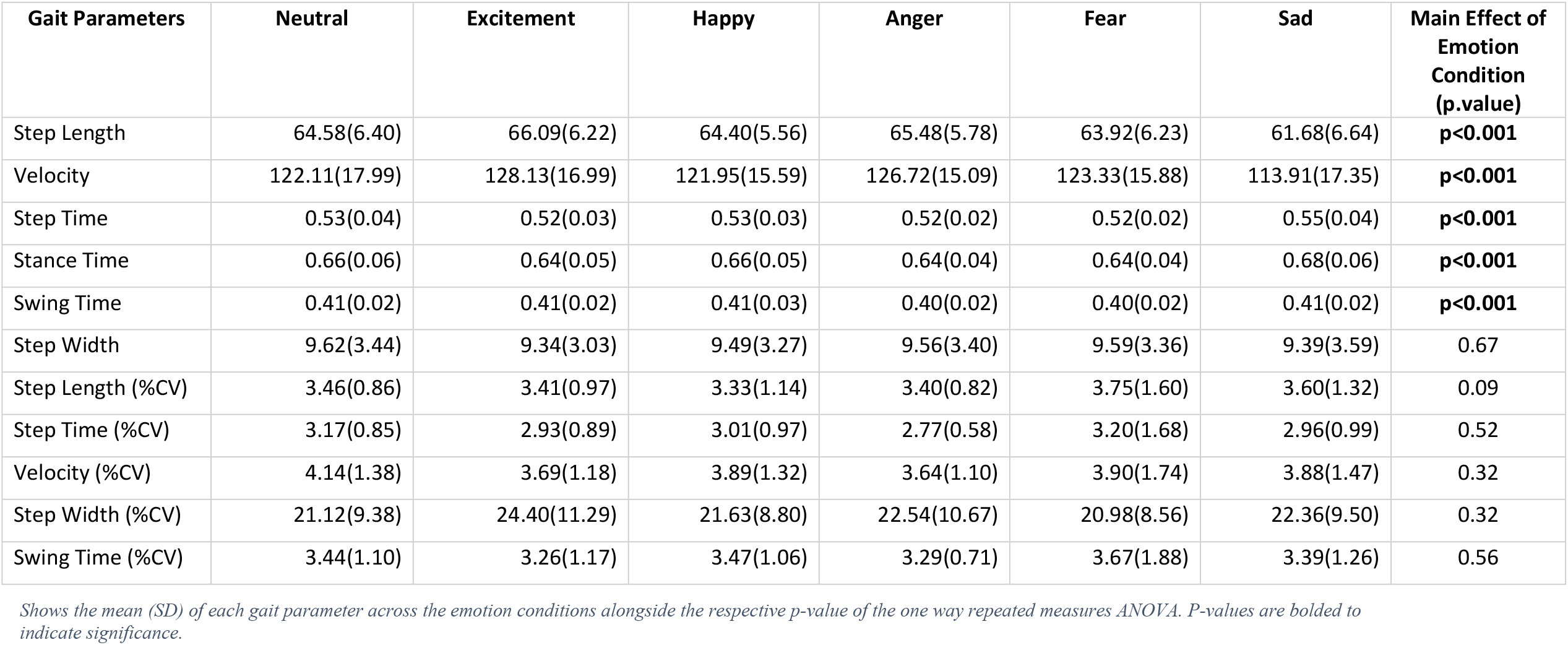
Summary results of gait parameters across emotional conditions.

### Pace Domain

A main effect of emotion condition was found for step length (χ^2^_Friedman_ (*df=5*) = 28.39, p<0.001, W_Kendall_ = 0.25) with a small effect size (Fig 1). Participants walked with smaller step length during the sadness condition compared to the other condition of anger, excitement, and the neutral condition. No other significant differences were observed between the other emotional conditions. Post hoc analyses showed that step length was significantly reduced during the sadness condition compared to the emotions of anger (p<0.001), excitement (p<0.001), and the neutral emotion (p=0.02).

**Fig 1.**
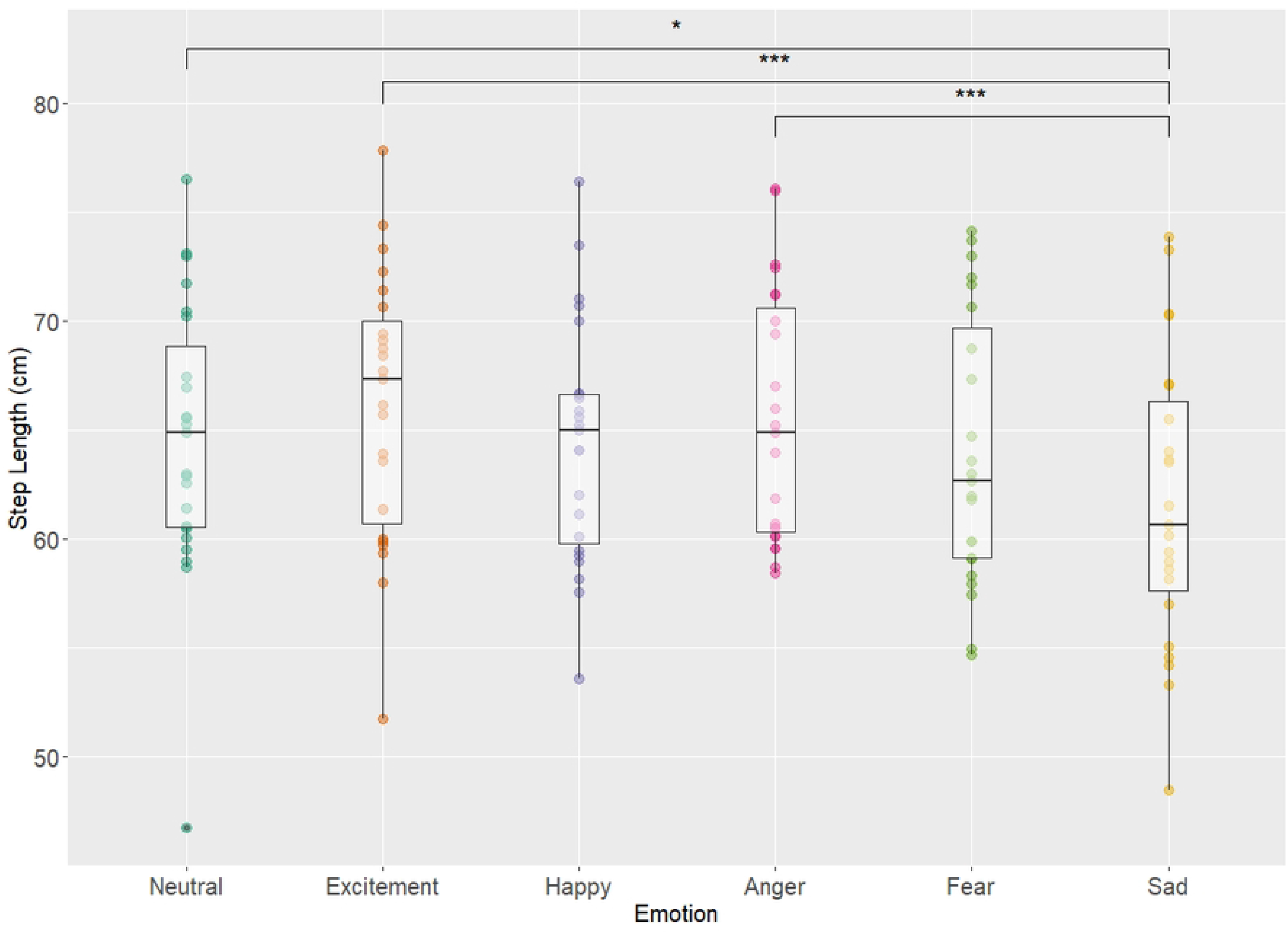
Step length and emotion. Displays the boxplot of each emotional condition and mean step length (cm) for each emotion alongside significant pairwise results. Main effect of emotion condition was found (χ^2^_Friedman_ (df=5) = 28.39, p =3.05e^-5^, W_Kendall_ = 0.25) with a small effect size. Adjusted p values are simplified as *, **, *** for p values <0.05, 0.01, 0.001 respectively.

A main effect of emotional condition was also found for gait velocity, (χ^2^ _Friedman_ (df = 5) =35.47, p<0.001, W_Kendall_ = 0.31) with a moderate effect size (Fig 2). Participants walked with the slowest gait velocity during the sadness condition compared to all other emotional conditions. No significant differences in gait velocity were observed when comparing the remaining emotional conditions. Post hoc analysis showed that gait velocity was significantly reduced during the sadness condition compared to the emotion conditions of excitement (p<0.001), anger (p<0.001), fear (p<0.001), neutral (p = 0.00493), and happiness (p = 0.0128).

**Fig 2.**
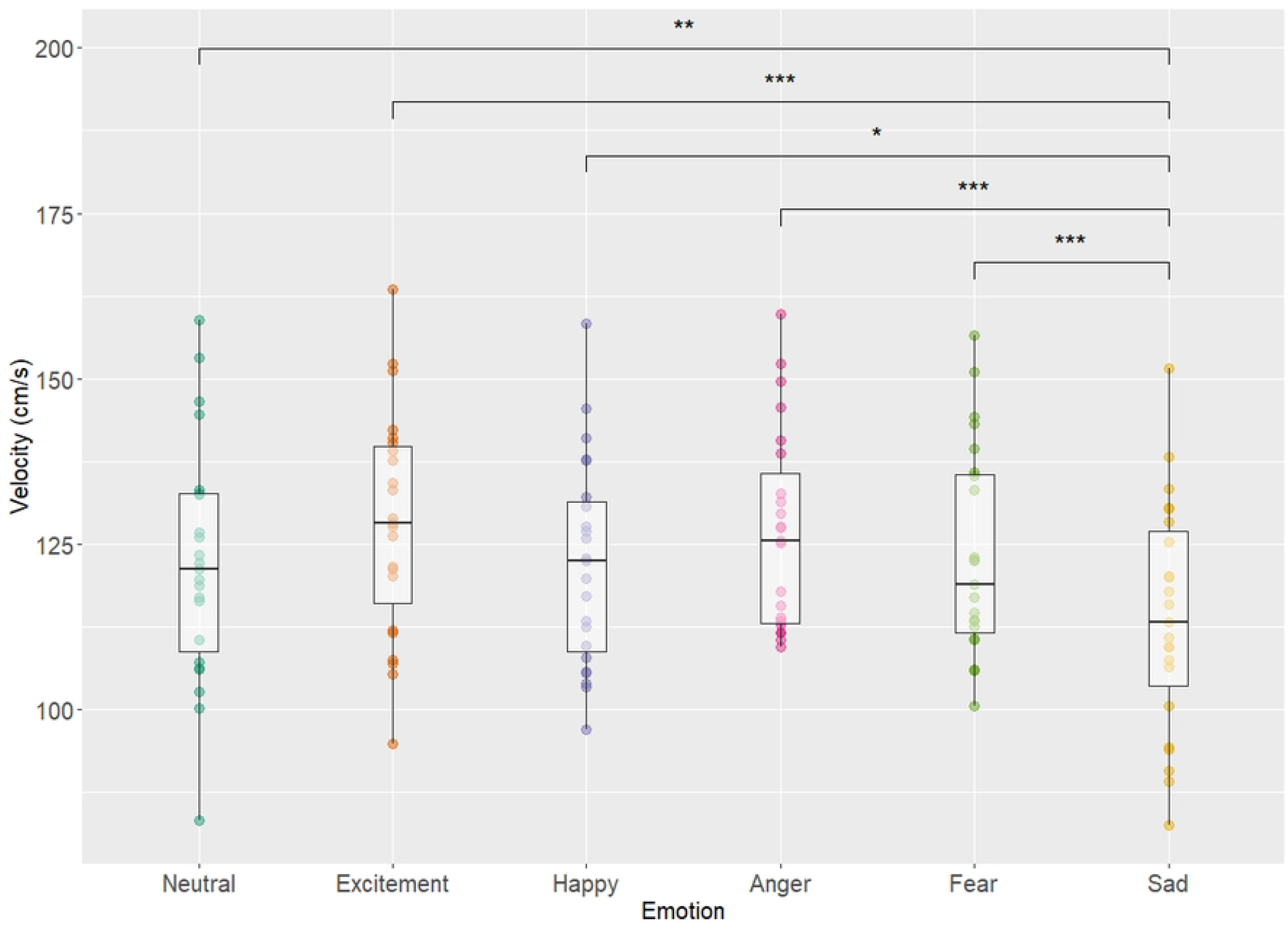
Gait velocity and emotion. Displays the boxplot of each emotional condition and velocity (cm/s) for each emotion alongside significant pairwise results. Main effect of emotional condition was found (χ^2^_Friedman_ (df = 5) =35.47, p=1.21e^-6^, W_Kendall_ = 0.31) with a moderate effect size. Adjusted p values are simplified as *, **, *** for p values <0.05, 0.01, 0.001 respectively.

### Rhythm Domain

Within the rhythm domain, all three gait outcomes (i.e., step time, stance time, and swing time) showed significant results. Fig 3(A) shows a main effect of emotional state condition was found for step time (χ^2^_Friedman_(df = 5) =37.75, p<0.001, W_Kendall_ = 0.33) with a moderate effect size (Fig3).A significant increase in step time was observed when comparing the condition of sadness to the conditions of fear, anger, and excitement but not the neutral condition. Significant decreases in step times were observed when comparing the excitement condition to the neutral and happiness conditions. Step time was also reduced during the fear condition compared to the conditions of excitement and happiness. Post hoc analysis revealed that participants had greater step time during the sadness condition compared to conditions of fear (p<0.001), anger (p<0.001), and excitement (p = 0.04). Participants walked with decreased step time during the excitement condition when compared to the neutral (p<0.001) and happiness (p=0.01) conditions. Finally, participants also walked with decreased step time during the fear condition compared to the conditions of happiness (p=0.04) and excitement (p=0.04).

**Fig 3.**
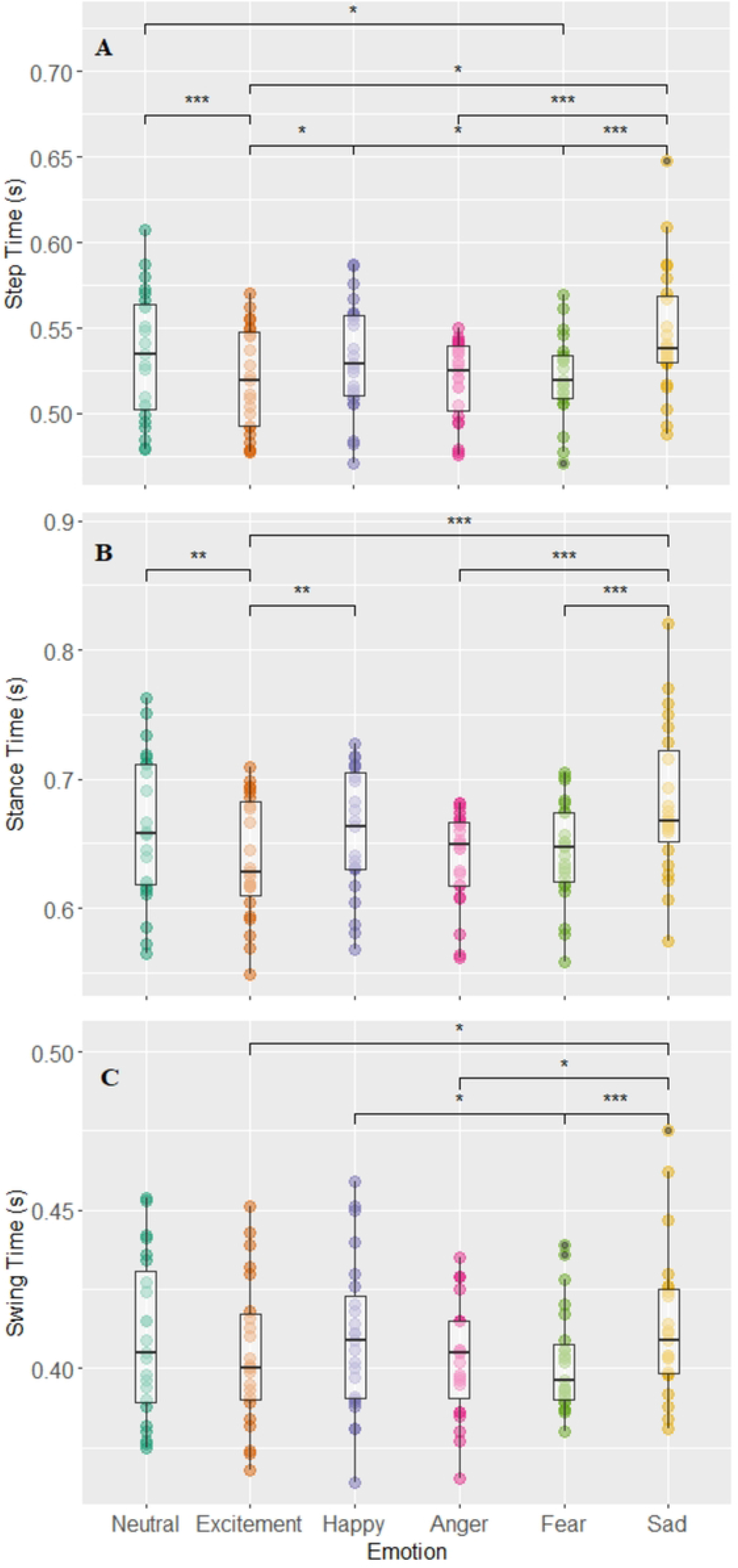
Step time, stance time, swing time, and emotion. Displays the boxplots of each emotional condition and the parameters of step time (A), stance time (B), and swing time (C). Main effects of emotion condition were found in all three parameters of step time χ^2^_Friedman_(df = 5) =37.75, p = 4.23e-7, W_Kendall_ = 0.33, stance time χ^2 Friedman^ (df = 5) =38.74, p = 2.88e-7, W_Kendall_ = 0.34, and swing time χ^2^_Friedman_ (df = 5) =23.77, p = 2.41e-4, W_Kendall_ = 0.21. Adjusted p values are simplified as *, **, *** for p values <0.05, 0.01, 0.001 respectively

A main effect of emotion condition was found for stance times (s) (χ^2^_Friedman_ (df = 5) =38.74, p<0.001, W_Kendall_ = 0.34), with a moderate effect size (Fig 3B). Results showed that participants walked with the longest stance times during the sadness condition and the shortest stance times during the excitement condition. Participants walked with increased stance time during sadness when compared to the conditions of fear, anger and excitement but not the neutral condition. Similar to step time, participants walked with a decreased stance time during the excitement condition when compared to the neutral and happiness condition. Post hoc analysis revealed the sadness condition had higher stance times than fear (p<0.001), anger (p<0.001), and excitement (p<0.001) conditions. The excitement condition also had lower stance time than that of neutral (p= 0.0083) and happy (p= 0.02) conditions.

A main effect of emotion condition was found for swing time (s) (χ^2^ _Friedman_ (df = 5) =23.77, p<0.001, W_Kendall_ = 0.21) with a small effect size (Fig 3C). Results show that participants walked with a greater swing time during the condition of sadness compared to the conditions of fear, anger, and excitement. A significant increase in swing time was also observed during the happiness condition when compared to the fear condition. No emotional condition showed any differences to the neutral condition when comparing swing time. Post hoc analysis revealed the sadness condition displayed longer swing times compared to fear (p<0.001), anger (p= 0.01), and excitement (p= 0.01) conditions. The fear condition also showed smaller swing time values compared to the happy condition (p= 0.01).

## Discussion

The primary objective of this study was to examine the influence of the emotional states of happiness, excitement, anger, fear, and sadness on spatiotemporal aspects of gait in a healthy adult non-actor population. It was hypothesized that emotions of happy, excitement, and anger would result in increases in gait speed when compared to the *neutral* condition while the opposite was expected for emotions of sadness and fear. This hypothesis was partly supported by the results, the sadness condition did show a decrease in gait speed compared to the neutral, confirming the hypothesis. However, the hypothesis was not supported in the other emotional state conditions, since gait speed was not different from the neutral condition when emotional states such as anger, fear, excitement or happiness were induced. It was also hypothesized that the condition of fear would result in an increase in gait variability compared to the neutral condition, however this was also not supported by the current results as changes to gait variability were not observed across any emotional state condition.

In the pace domain, sadness, above all other emotion conditions showed a large discrepancy in gait parameters such as reductions in step length and gait velocity which is in concordance with previous literature. Barliya et al. showed that, the sadness condition resulted in the slowest gait speed and that fear, happy, and anger conditions showed faster gait compared to sadness, which is reflected in the results of this study [10]. Results also show the condition of sadness had the shortest average step length compared to the other conditions which is supported by previous literature [9]. Current results show that decrease in parameters of gait speed in sadness were similar to decreases in gait speed when comparing healthy controls and patients with major clinical depression. Lemke et al. [6] showed that patients with major depression had a reduction of ∼0.23m/s in gait velocity compared to healthy controls. This study showed that sadness condition resulted in difference of ∼0.08m/s when compared to the neutral condition. This difference is greater when comparing excitement’s velocity of 1.28 m/s to sadness’ 1.13 m/s which results in a difference of ∼0.15 m/s which is closer to the gait speed reduction seen in healthy controls vs patients with clinical depression. While it is not accurate to attempt to equate sadness with major depression, it is evident that emotions can have the capacity to meaningfully change gait speed. Therefore, the emotion of sadness (as it pertains to gait speed and step length) perhaps could be considered a potential confound for future gait analysis and research in healthy and clinical populations.

In the rhythm domain, the sadness condition was also different from other emotional conditions (higher stance time and step time compared to excitement, anger, fear) however none of the variables showed a difference between the neutral condition and sadness. These results also are supported by previous literature, as Barliya et al. [10] showed that the sadness condition occupied a lower % swing phase duration and thus the inverse is true where a larger % stance phase duration is observed. Differences in gait between the neutral condition and the conditions of excitement and fear were observed in step time (both excitement and fear conditions showed decreased step times compared to neutral) and stance time (excitement condition lower than neutral).

Thus, with these results it is important to highlight that the impact of emotional states on gait may not just be a general modulator of pace and overall gait speed. Compared to the neutral condition, the conditions of fear and excitement, did not show changes in the pace domain but did show changes in the rhythm domain. It could be possible that emotional states, especially those of fear and excitement may impact stepping characteristics and the rhythm of gait, regardless of pace/speed. Future work could possibly investigate this discrepancy between the two domains and emotions further to understand how one domain may be affected but not the other.

Based on findings from the current study, emotional states did not influence gait variability, nor postural control. Previous research showed that different mood disorders, namely increased anxiety in older adults with PD showed greater step-to-step variability [19] and depression was shown to impact swing time variability (although this exists in the pace domain according to the 5-factor model of gait) [12]. The sample in this study involved young healthy adults. It is possible that young healthy adults maintain a high degree of automaticity over their gait regardless of emotional conditions leading to fewer changes in their step-to-step variability. Studies that have investigated dual tasking have shown that an increased cognitive load and cognitive demand consistently have a destabilizing effect on gait on older adults and thus impacting gait variability [20]. Perhaps the intensity of emotions elicited in this study were not sufficient enough of a cognitive demand to cause changes in gait variability in young healthy adults.

## Limitations, and Future Directions

In terms of emotion elicitation, while VR and video was a valid way to elicit emotions, the videos remain still somewhat subjective and are prone to resulting in different interpretations causing inter-participant discrepancy in the SAM reporting and their own self-reported emotions. The usage of the 23-person emotional dataset was an attempt to address this discrepancy in emotional response and reduce the inter-participant variability in report of SAMs and emotional reports. Future studies should consider including a physiological measure of some of these dimensions such as electrodermal skin conductance (EDA) to measure arousal levels, which could give a better indication of the effectiveness of emotion elicitation.

It remains unclear whether the results of this study regarding emotional states and unchanging gait variability parameters would be the same for older adults or a clinical population. Older adults, and patients with depression may walk with increased rumination, resulting in increased cognitive load, and increased distractors as they walk—all which culminates in a modified gait pattern [21,22]. This could potentially introduce more avenues for affective, particularly negative affective states like sadness to further alter gait behavior in the population. Further work is needed to study how emotional states affects older adults and those in clinical populations.

## Conclusion

The results from this current study show that the emotions of sadness and excitement affect gait in young healthy adults. The main findings show that sadness resulted in smaller steps, reduced gait speed, increased step time and stance times whilst the opposite was observed during excitement. Emotions main impacted gait parameters within the pace and rhythm domain, but less so for aspects of gait variability and postural control. It is recommended that future research in both clinical and experimental settings should possibly consider evaluating the emotional state of an individual, particularly the emotions of sadness and excitement as it showed to have influenced young healthy adult gait.

## Acknowledgements

We would like to acknowledge all the assistance, encouragement, and support of the NeuroCognition and Mobility Lab team which helped make this study possible. This work was supported by an NSERC Discovery Grant (KEM).

## References

1. Deligianni F, Guo Y, Yang GZ. From Emotions to Mood Disorders: A Survey on Gait Analysis Methodology. IEEE J Biomed Health Inform. 2019;23(6):2302–16.

2. Takakusaki K. Neurophysiology of gait: From the spinal cord to the frontal lobe. Movement Disorders. 2013;28(11):1483–91.

3. Radovanović S, Jovičić M, Marić NP, Kostić V. Gait characteristics in patients with major depression performing cognitive and motor tasks while walking. Psychiatry Res [Internet]. 2014 Jun x30 [cited 2023 Feb 27];217(1–2):39–46. Available from: https://pubmed.ncbi.nlm.nih.gov/24613201/

4. Sanders RD, Gillig PM. Gait and its assessment in psychiatry. Psychiatry (Edgmont) [Internet]. 2010 Jul [cited 2023 Feb 27];7(7):38. Available from: /pmc/articles/PMC2922365/

5. Michalak J, Troje NF, Fischer J, Vollmar P, Heidenreich T, Schulte D. Embodiment of sadness and depression--gait patterns associated with dysphoric mood. Psychosom Med [Internet]. 2009 Jun [cited 2023 Feb 27];71(5):580–7. Available from: https://pubmed.ncbi.nlm.nih.gov/19414617/

6. Lemke MR, Wendorff T, Mieth B, Buhl K, Linnemann M. Spatiotemporal gait patterns during over ground locomotion in major depression compared with healthy controls. J Psychiatr Res. 2000;34(4–5):277–83.

7. Ehgoetz Martens KA, Ellard CG, Almeida QJ. Does anxiety cause freezing of gait in Parkinson’s disease? PLoS One. 2014;9(9).

8. Roether CL, Omlor L, Christensen A, Giese MA. Critical features for the perception of emotion from gait. J Vis. 2009;9(6):1–32.

9. Halovic S, Kroos C. Not all is noticed: Kinematic cues of emotion-specific gait. Hum Mov Sci. 2018;57(July 2017):478–88.

10. Barliya A, Omlor L, Giese MA, Berthoz A, Flash T. Expression of emotion in the kinematics of locomotion. Exp Brain Res. 2013;225(2):159–76.

11. Lord S, Galna B, Verghese J, Coleman S, Burn D, Rochester L. Independent domains of gait in older adults and associated motor and nonmotor attributes: Validation of a factor analysis approach. Journals of Gerontology -Series A Biological Sciences and Medical Sciences. 2013;68(7):820–7.

12. Dragašević-Mišković NT, Bobić V, Kostić M, Stanković I, Radovanović S, Dimitrijević K, et al. Impact of depression on gait variability in Parkinson’s disease. Clin Neurol Neurosurg. 2021;200(February 2020).

13. Li BJ, Bailenson JN, Pines A, Greenleaf WJ, Williams LM. A public database of immersive VR videos with corresponding ratings of arousal, valence, and correlations between head movements and self report measures. Front Psychol. 2017;8.

14. Marín-Morales J, Higuera-Trujillo JL, Greco A, Guixeres J, Llinares C, Gentili C, et al. Real vs. immersive-virtual emotional experience: Analysis of psycho-physiological patterns in a free exploration of an art museum. PLoS One. 2019;14(10):1–24.

15. Lorenzetti V, Melo B, Basílio R, Suo C, Yücel M, Tierra-Criollo CJ, et al. Emotion regulation using virtual environments and real-time fMRI neurofeedback. Front Neurol. 2018;9:1–15.

16. Higuera-Trujillo JL, López-Tarruella Maldonado J, Llinares Millán C. Psychological and physiological human responses to simulated and real environments: A comparison between Photographs, 360° Panoramas, and Virtual Reality. Appl Ergon. 2017; 65:398–409.

17. Perera S, Smith C, Coffman L, Brach J. Number of Steps Needed for Reliable Gait Variability Measurement. Gerontologist [Internet]. 2016 Nov 1 [cited 2021 Oct 31];56(Suppl_3):335–6. Available from: https://academic.oup.com/gerontologist/article/56/Suppl_3/335/2574681

18. Patil I. Visualizations with statistical details: The “ggstatsplot” approach. J Open Source Softw [Internet]. 2021 May 25;6(61):3167. Available from: https://joss.theoj.org/papers/10.21105/joss.03167

19. Ehgoetz Martens KA, Silveira CRA, Intzandt BN, Almeida QJ. Overload from anxiety: A non-motor cause for gait impairments in Parkinson’s disease. Journal of Neuropsychiatry and Clinical Neurosciences. 2018;30(1):77–80.

20. Hollman JH, Kovash FM, Kubik JJ, Linbo RA. Age-related differences in spatiotemporal markers of gait stability during dual task walking. Gait Posture. 2007 Jun 1;26(1):113–9.

21. Thomsen DK. The association between rumination and negative affect: A review. Vol. 20, Cognition and Emotion. 2006. p. 1216–35.

22. Iqbal N, Dar KA. Negative affectivity, depression, and anxiety: Does rumination mediate the links? J Affect Disord [Internet]. 2015 Aug 1 [cited 2023 Mar 19]; 181:18–23. Available from: https://pubmed.ncbi.nlm.nih.gov/25913918/

